# Genome sequencing and *de novo* and reference-based genome assemblies of *Bos indicus* breeds

**DOI:** 10.1101/2022.08.27.505546

**Authors:** Abhisek Chakraborty, Manohar S. Bisht, Rituja Saxena, Shruti Mahajan, Joby Pulikkan, Vineet K. Sharma

## Abstract

*Bos indicus* is a domestic cattle species with many indigenous breeds found in India and is important for dairy, draught work, and other household activities. Distinct phenotypic differences are observed among the breeds of this species; however, their whole genome sequences were not available. Therefore, in this study, we performed the whole genome sequencing using Illumina short-read technology to construct draft genome assemblies of four *B. indicus* breeds; Ongole, Kasargod Dwarf, Kasargod Kapila, and Vechur, of which Vechur is known as the smallest cow of the world. We also report the first *de novo* genome assemblies of these native *B. indicus* breeds. Further, we constructed the 18S rRNA marker gene sequences of these *B. indicus* breeds, which were not yet known. Genomic analysis helped to identify the distinct bovine phenotypic characteristics-related and other biological process-related genes in this species compared to *B. taurus*, and to gain comparative genomic insights between the dwarf and non-dwarf breeds of this species.

## INTRODUCTION

Livestock plays a key role in maintaining the economy of both developed and developing countries. In India, this provides livelihood support to more than eighty million rural households engaged in dairying, and thus making India a global leader among the dairying nations [1]. So far, 50 distinct *Bos indicus* cattle breeds (https://nbagr.icar.gov.in/en/registered-cattle/) have been reported from India, which have evolved by adapting to various agroclimatic conditions of India pertaining to their survival in harsh climatic conditions, ability to perform on poor quality feed and fodder, and having innate immunity to various diseases (Sharma et al., 2015; Pramod et al., 2018; Shivakumara et al., 2018). By contrast, hump-less *Bos taurus* are less adaptive to heat stress and to temperate climatic conditions (Mwai et al., 2015; Cooke et al., 2020). Furthermore, the Indicine cattle breeds produce milk rich in A2 casein variant, which is better for health (Mishra et al., 2009), whereas the *B. taurus* breeds are generally carriers of A1 casein variant (Kamiński et al., 2007; Patel et al., 2020).

Cattles in Indian subcontinent are also known as zebu cattle (*Bos primigenius indicus*) that evolved along with taurine cattle (*Bos primigenius taurus*) from extinct wild aurochs (*Bos primigenius*) nearly between 250,000 and 330,000 years ago, into two distinct lineages (Pitt et al., 2019; Fernandes Júnior et al., 2020). Previous studies have identified the diverse breed of zebu cattle (*B. indicus*) present in India (Dixit et al., 2020; Dixit et al., 2021). However, the comprehensive genomic and genetic analysis of various indigenous cow breeds of India is yet to be carried out.

The 50 registered breeds of *B. indicus* in India have been categorized as pure milking breeds (Deoni, Gir, Red Sindhi, Sahiwal, and Tharparkar), dual purpose, and triple purpose breeds [1,2]. However, several Indicine cattle are yet to be described as a breed, for example, Kasargod Dwarf and its variety Kasargod Kapila are one of the non-descriptive indigenous dwarf cattle. However, in this study and in the following text, we have referred both (Kasargod Dwarf and its variety Kasargod Kapila) as breeds for the ease of writing and readability. Kasargod Dwarf derives its name from the mountain ranges of Kasaragod, Kerala, India. The average body length, height and weight of an adult Kasargod cattle ranges from 101 to 106 cm, 97 to 102 cm, and 164 to 178 kg, respectively (Iype et al., 2016). Despite being short in stature, Kasargod Dwarf possesses various characteristics that distinguish them from exotic cattle e.g., high nutritional quality of milk, high disease and heat tolerance, and increased feed efficiency (Deepthi et al., 2021). Another important and interesting breed is known as Vechur cow, the smallest described cattle breed of the world that derives its name from a village “Vechoor” in Kottayam district of Kerala. For adult Vechur cattle, average body height, length, and weight are recorded in ranges from 84 to 99 cm, 93 to 104 cm, and 95 to 178 kg, respectively (Radhika et al., 2018). It is known for its higher milk production capacity (2–3 litres per day) given its dwarf size and compared to the other known non-descriptive cow breeds. Vechur cattle are also well-known for their resistance to viral, bacterial, and parasitic diseases compared to the exotic cattle and their crossbreds (Shivakumara et al., 2018).

Ongoles are a very majestic large-sized humped and triple purpose (draught, milk and meat animals) cattle breed (Revista and 1993). Like other Indian breeds of cattle, Ongoles also got their name from their main breeding city Ongole from the Indian state of Andhra Pradesh. However, Ongole cattle is an Indian breed, but it has been internationally known due to the development of exotic breeds like ‘Santa Getrudis’, ‘American Brahman’ etc. and is used widely for production of beef in Latin American countries (Beja-Pereira et al., 2003; Metta et al., 2004). Average body length, height, and weight of an adult Ongole are recorded as 141.83 cm, 145.7cm and 408 kg, respectively (Animal Genetic Resource of India; http://14.139.252.116/agris/bridDescription.aspx).

The previous genomic studies of the Indian cow breeds were focused on a few characteristics such as heat tolerance, stature, and physiological characteristics like milk type A1/A2, disease resistance, etc. (Mishra et al., 2009; Pryce et al., 2011; Elayadeth-Meethal et al., 2021). However, the absence of genome sequence of these unique Indicine cow breeds is a limiting factor in elucidating the genomic basis of origin of these distinctive phenotypic characters. Therefore, in this study, we performed the genome sequencing of *B. indicus* species - Kasargod Kapila, Kasargod Dwarf, Vechur and Ongole breeds using Illumina short-read sequencing platform and performed *de novo* and reference-based genome assemblies followed by genomic analysis. Although a reference-based genome assembly of *B. indicus* cattle was once performed in a previous study (Canavez et al., 2012), the *de novo* genome assembly of this species is not yet available. Therefore, in this study, we also carried out the first draft *de novo* genome assembly of *B. indicus* breeds (Ongole, Vechur, Kasargod Dwarf, and Kasargod Kapila), and analyzed the structures of the genes responsible for the distinct phenotypic characteristics of these cattle breeds. Further, this study also reports the first draft sequences of the 18S rRNA marker gene sequences of these four *B. indicus* breeds.

## MATERIALS AND METHODS

### Sample collection, library preparation, and sequencing

The blood samples were collected and transported on dry ice to the laboratory at IISER Bhopal and were stored immediately at −80°C till further processing. Genomic DNA was extracted from the blood samples using DNeasy Blood and Tissue kit (Qiagen, CA, USA). The extracted DNA were quantified on Qubit 2.0 fluorometer using qubit dsDNA BR assay kit (Invitrogen, USA). The quantified DNA was used for multiple library preparation using Nextera XT (Illumina Inc., USA), TruSeq DNA PCR free (Illumina Inc., USA) and Collibri PCR free PS DNA (ThermoFisher, United states) library preparation kits. Quantification of the libraries were performed on Qubit 2.0 fluorometer using qubit dsDNA HS assay kit (Invitrogen, USA) and qPCR using KAPA SYBR FAST qPCR master mix with primer premix and Illumina standards (KAPA Biosystems, United States). The library size was estimated on Agilent Bioanalyzer 2100 using high sensitivity DNA kit (Agilent, United States). The normalized libraries were sequenced for 150 bp paired-end reads on Illumina NextSeq 500 platform using NextSeq 500/550 v2 and v2.5 sequencing reagent kits (Illumina Inc., USA) at NGS facility, IISER Bhopal. For Ongole DNA sample, amplification of the whole genome was performed using Genomiphi V2 DNA amplification kit (GE Healthcare, UK) and purified using Ampure XP magnetic beads (Beckman coulter, USA). The amplified DNA was used to prepare library using TruSeq DNA Nano library preparation kit (Illumina

Inc., USA). The library was quantified on Qubit 4.0 fluorometer using qubit dsDNA HS assay kit (Invitrogen, USA). The library quality was assessed on Tapestation 4150 (Agilent, United states) using high sensitivity D1000 screentape. The normalized library was loaded on Illumina sequencing platform for generating 150 bp paired-end reads.

### Genomic data pre-processing

Raw data were demultiplexed using Bcl2fastq, and the obtained FastQ files containing demultiplexed reads were trimmed and filtered using Trimmomatic v0.39 (Bolger et al., 2014). Output files containing unpaired reads (where one of the matching paired-end read passed quality parameter) were concatenated into a single file. Reads were filtered for a minimum length of 40 bp and 2 mismatches to be allowed in the seed matching with seed length of 16, palindrome clip threshold of 30, and simple clip threshold of 10. The low-quality bases or N’s were removed from the leading and trailing ends of the reads with the quality threshold of 15. The reads were scanned with a sliding window of 15 bp, and the reads were trimmed when the average PHRED quality score per base went below 15.

### Reference-based assembly and variant calling

Quality-filtered reads (both paired-end and unpaired) were mapped to the reference *Bos taurus* genome (ARS-UCD 1.2) obtained from Ensembl genome browser 104 (Hubbard et al., 2002) using Bowtie2 v2.4.1 (Langmead and Salzberg, 2012). Conversion of the obtained SAM files to BAM format was performed using SAMtools v1.13 “view” (Li et al., 2009). BAM files were sorted based on their coordinates, and duplicates were marked using Picard (v2.26.6) “MarkDuplicates” (https://broadinstitute.github.io/picard/). Variant calling in the coding genes was performed using BCFtools (v1.14) “mpileup” (Narasimhan et al., 2016), and further filtering of false-positive variants was carried out based on the following parameters - variant sites with quality ≥50, sequencing depth ≥50. Consensus reference-based genome assemblies for each breed were obtained using BCFtools (v1.14) “consensus”

(Narasimhan et al., 2016). Quality-filtered paired-end and unpaired reads were mapped onto the respective reference-based genome assemblies of each breed using Bowtie2 v2.4.1, and the mapping statistics were calculated using SAMtools (v1.13) “flagstat”. Further, percentage sequence variation and the number of scaffolds showing sequence variation were calculated from the obtained filtered VCF files. BLASTN was used to check the sequence homology against the *B. taurus* coding gene sequences with 97% identity threshold, and e-value of 10^−9^.

### *de novo* assembly of Indicine breed

The quality-filtered paired-end reads of all four breeds of *B. indicus* were *de novo* assembled individually using SPAdes v3.15.3 (Bankevich et al., 2012) with k-mer value of 99 and “only-assembler” option. Further scaffolding of these *de novo* assemblies were generated by mapping onto *B. taurus* genome assembly using Satsuma2 “Choromosemble” (Grabherr et al., 2010) module. 15 *B. indicus*-specific gene sequences related to bovine characteristics such as milk quality, stature, metabolism and immune response (Pryce et al., 2011; Dixit et al., 2020; Dixit et al., 2021; Elayadeth-Meethal et al., 2021) were obtained from NCBI/UniProt database, and were mapped onto the genome assemblies of *B. indicus* breed Ongole using Exonerate v2.2.0 (https://github.com/nathanweeks/exonerate) to check their presence and find the exon-intron structures. The same genes specific to *B. taurus* species were also mapped to *B. taurus* reference genome (ARS-UCD 1.2) obtained from Ensembl genome browser 104 to compare the difference.

Jellyfish v2.2.10 (Marçais and Kingsford, 2011) was used to construct a k-mer frequency distribution using the quality filtered paired-end reads of Ongole. Percent heterozygosity of *B. indicus* genome was calculated with this k-mer frequency distribution using GenomeScope v2.0 (Ranallo-Benavidez et al., 2020).

The detailed computational analysis workflow is explained in **Figure 1**.

**Figure 1.**
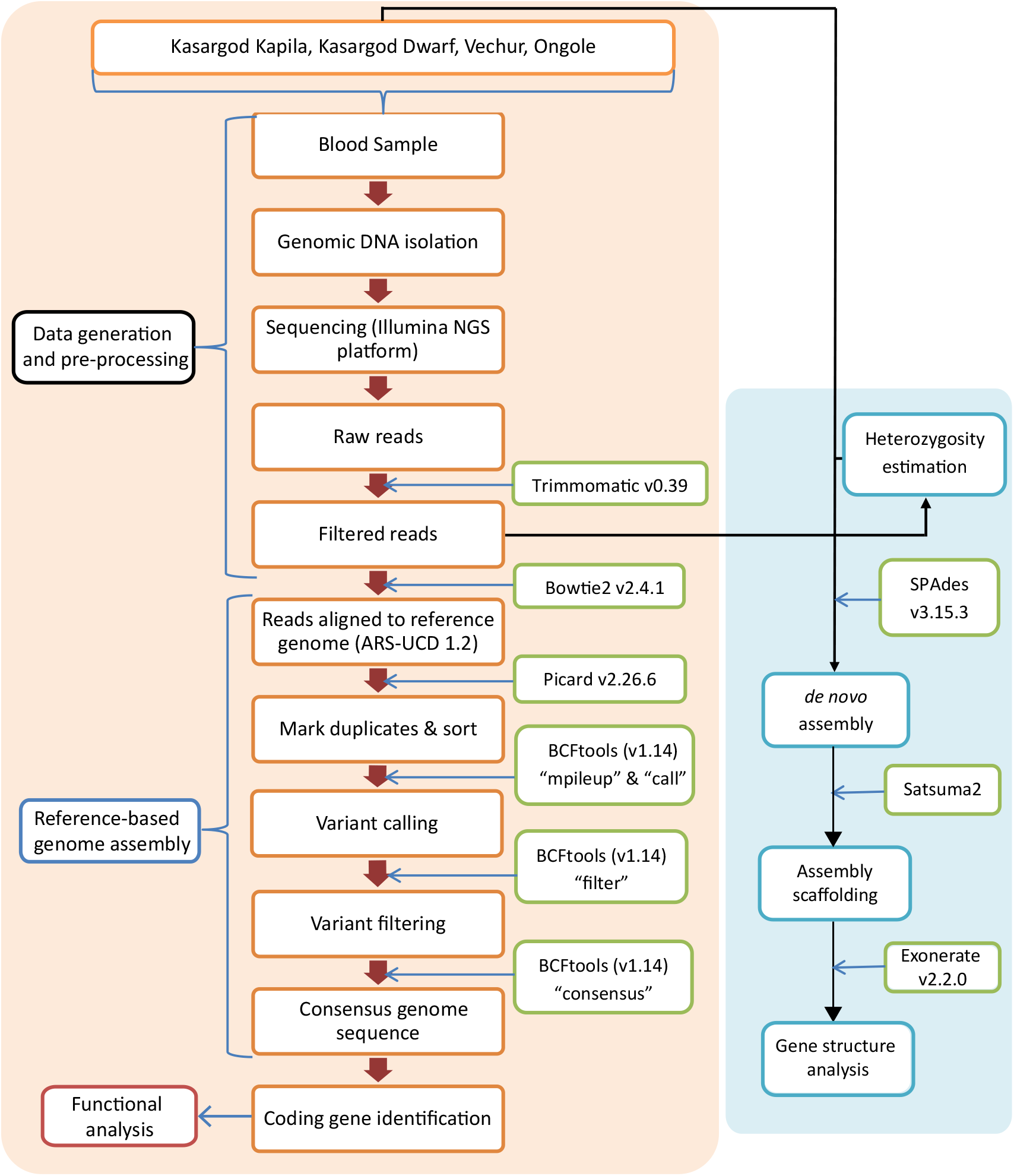
The complete genomic data analysis workflow for *B. indicus* breeds.

### Species identification with marker genes

The extracted DNA of Ongole breed was used for amplifying Cytochrome c oxidase subunit I (COI) and 18S rRNA marker genes for species identification. 18S rRNA of length ~1.8 Kb was amplified in three fragments, which were further assembled to form a complete 18S rRNA gene. All amplifications were performed using Taq Polymerase (Invitrogen, United States). Additives, i.e., Dimethyl sulphonate (DMSO) and Bovine serum albumin (BSA) were used in amplifying first and second fragments of 18S rRNA gene. The amplified products were checked on 2% agarose electrophoresis and purified using Purelink PCR purification kit (Invitrogen, United States). The purified amplicons were quantified on Qubit 2.0 using qubit dsDNA BR assay kit (Invitrogen, United States). 40 ng of amplified products were used for Sanger sequencing at the central instrumentation facility at IISER Bhopal.

## RESULTS

### Genome sequencing

Sequencing using Illumina NGS platform generated raw data of 141.1 Gb, 49.9 Gb, 47.5 Gb, and 40.3 Gb, corresponding to the coverage of 52.2x, 18.5x, 17.6x, and 14.9x for Ongole, Kasargod Dwarf, Kasargod Kapila, and Vechur, respectively (estimated genome size ~2.7 Gbp https://pag.confex.com/pag/xxv/webprogram/Paper26680.html). After trimming and quality-filtering using Trimmomatic, 132.46, 46.8 Gb, 44.5 Gb, and 39.1 Gb data was obtained for Ongole, Kasargod Dwarf, Kasargod Kapila, and Vechur, respectively.

### Reference-based genome assembly

The consensus reference-based genome assemblies for the selected breeds of *B. indicus* were constructed by mapping the quality-filtered reads on *B. taurus* genome (ARS-UCD 1.2) using Bowtie2 v2.4.1 (Langmead and Salzberg, 2012). Consensus genome sequences for each breed were obtained using BCFtools “consensus” (Narasimhan et al., 2016) (N50 ~ 103.33 Mbp). Mapping percentage for each consensus genome assembly, and number of scaffolds containing variation are mentioned in **Table 1**.

**Table 1.** Summary statistics for reference-based genome assembly

| Sample name | Mapping percentage<br>(%) | Number of scaffolds<br>containing variation |
| --- | --- | --- |
| Kasargod Kapila | 89.31 | 1,500 |
| Kasargod Dwarf | 90.24 | 1,629 |
| Vechur | 86.32 | 814 |
| Ongole | 89.07 | 1,024 |

BLASTN alignment against the coding gene set of *B. taurus* (37,538 genes) identified a total of 36,636, 36,633, 35,993, and 36,639 genes with 100% sequence identity in Kasargod Kapila, Kasargod Dwarf, Ongole, and Vechur breeds, respectively. A total of 26 genes were found in the above breeds of *B. indicus* that were less identical to the *B. taurus* genes (**Table 2**). Additionally, five more genes were identified that could be mapped in the dwarf breeds of *B. indicus* but were not found in Ongole (with stringent BLASTN criteria). Similarly, one gene was mapped in Ongole, but could not be mapped in any dwarf breed of *B. indicus*.

**Table 2.** List of genes that showed a difference in sequence identity between *B. taurus* and *B. indicus*, and between dwarf and non-dwarf breeds of *B. indicus*

| **Genes that showed low sequence identity in *B. indicus*** |  |  |
| --- | --- | --- |
| **Ensembl gene IDs** | **Gene name** | **Gene description/biological function** |
| ENSBTAT00000036708.5 | Novel gene | Chromatin binding, histone binding |
| ENSBTAT00000036961.4 | Novel gene | Structural constituent of ribosome |
| ENSBTAT00000063969.2 | OR51A4* | Olfactory receptor family 51 subfamily A member 4 |
| ENSBTAT00000066353.1 | Novel gene | Keratinization, epidermis formation |
| ENSBTAT00000067159.1 | Novel gene | - |
| ENSBTAT00000069026.1 | OLFR1045* | Olfactory receptor 1045 |
| ENSBTAT00000069107.1 | Novel gene | Extracellular matrix structural constituent |
| ENSBTAT00000069874.1 | ZNF333* | Zinc finger protein 333 |
| ENSBTAT00000071298.1 | Lce3b* | Late cornified envelope 3C |
| ENSBTAT00000073804.1 | RPS19* | 40S ribosomal protein S19 |
| ENSBTAT00000074029.1 | TRBV21* | T cell receptor beta, variable 21 |
| ENSBTAT00000075233.1 | Novel gene | Olfactory receptor activity |
| ENSBTAT00000076752.1 | Novel gene | - |
| ENSBTAT00000076772.1 | LHFPL3* | LHFPL tetraspan subfamily member 3 |
| ENSBTAT00000076797.1 | Novel gene | - |
| ENSBTAT00000077152.1 | Novel gene | Keratinization, epidermis formation |
| ENSBTAT00000078401.1 | IGKV* | Immunoglobulin kappa chain variable |
| ENSBTAT00000078580.1 | OR4L17 | Olfactory receptor family 4 subfamily L<br>member 17 |
| ENSBTAT00000079019.1 | LCE1B | Late cornified envelope 1B |
| ENSBTAT00000080177.1 | Novel gene | Olfactory receptor 2H2-like |
| ENSBTAT00000080901.1 | CFAP299* | Cilia and flagella associated protein 299 |
| ENSBTAT00000081050.1 | OR7E200 | Olfactory receptor family 7 subfamily E<br>member 200 |
| ENSBTAT00000081366.1 | Novel gene | - |
| ENSBTAT00000084909.1 | MYADM* | Myeloid-associated differentiation marker |
| ENSBTAT00000085337.1 | ZNF71 | Zinc finger protein 71 |
| ENSBTAT00000086268.1 | Novel gene | Immune response |
| **Genes that showed low sequence identity in dwarf breeds compared to Ongole** |  |  |
| ENSBTAT00000020097.6 | OR52N4I | Olfactory receptor family 52 subfamily N<br>member 4I |
| ENSBTAT00000064254.2 | OR2O2 | Olfactory receptor family 2 subfamily O<br>member 2 |
| ENSBTAT00000066609.1 | UBE2D3 | Ubiquitin conjugating enzyme E2 D3 |
| ENSBTAT00000084042.1 | Novel gene | Response to bacterial infection |
| ENSBTAT00000084273.1 | TRAV8-7* | T cell receptor alpha variable 8-7 |
| **Gene that showed low sequence identity in Ongole compared to dwarf breeds** |  |  |
| ENSBTAT00000080177.1 | OR2H4* | Olfactory receptor family 2 subfamily H<br>member 4 |

\*Gene names were derived from orthologous genes in other species from Ensembl database

### *de novo* assembly of Indicine breeds

The draft *de novo* genome assemblies for all the *B. indicus* breeds had the sizes ranging between 1.98 - 3.42 Gbp (**Table 3**). *de novo* genome assemblies of Ongole, Kasargod Dwarf, Kasargod Kapila, and Vechur constructed using SPAdes and Satsuma2 resulted in N50 values of 97 Mbp, 72 Mbp, 73 Mbp, and 96 Mbp, respectively (**Table 3**). The percent heterozygosity was estimated to be 0.927% for *B. indicus* species (Ongole breed) (**Figure 2**). **18S rRNA marker gene construction**

**Table 3.** Summary statistics of the *de novo* genome assemblies of *B. indicus* breeds

| Parameters | Name of *B. indicus* breeds |  |  |  |
| --- | --- | --- | --- | --- |
|  | Ongole | Kasargod Dwarf | Kasargod Kapila | Vechur |
|  | Value | Value | Value | Value |
| Contigs ($\geq 0$ bp) | 3,763,658 | 532,785 | 437,825 | 382,190 |
| Contigs ($\geq 1$ Kbp) | 4,684 | 811 | 641 | 3,216 |
| Contigs ($\geq 5$ Kbp) | 366 | 253 | 215 | 215 |
| Contigs ($\geq 10$ Kbp) | 236 | 166 | 165 | 155 |
| Contigs ($\geq 25$ Kbp) | 144 | 104 | 100 | 104 |
| Contigs ($\geq 50$ Kbp) | 97 | 69 | 71 | 72 |
| Total length (bp)<br>($\geq 0$ bp) | 3,418,321,891 | 2,058,585,533 | 1,978,806,794 | 2,573,575,944 |
| Total length (bp)<br>($\geq 1$ Kbp) | 2,883,554,001 | 1,912,054,931 | 1,847,133,685 | 2,452,655,955 |
| Total length (bp)<br>($\geq 5$ Kbp) | 2,876,941,510 | 1,910,973,226 | 1,846,284,924 | 2,448,608,000 |
| Total length (bp)<br>($\geq 10$ Kbp) | 2,876,076,773 | 1,910,357,025 | 1,845,935,974 | 2,448,179,362 |
| Total length (bp)<br>($\geq 25$ Kbp) | 2,874,592,721 | 1,909,389,855 | 1,844,955,485 | 2,447,434,916 |
| Total length (bp)<br>($\geq 50$ Kbp) | 2,872,922,794 | 1,908,177,327 | 1,843,977,665 | 2,446,327,213 |
| Contigs | 3,763,658 | 532,785 | 437,825 | 382,190 |
| Largest contig (bp) | 173,062,328 | 114,406,614 | 111,759,797 | 148,780,455 |
| Total length (bp) | 3,418,321,891 | 2,058,585,533 | 1,978,806,794 | 2,573,575,944 |
| GC (%) | 42.08 | 42.20 | 40.28 | 39.96 |
| N50 (bp) | 97,009,571 | 71,682,961 | 72,991,627 | 96,252,292 |
| N75 (bp) | 59,021,997 | 48,612,102 | 44,589,875 | 64,286,014 |
| L50 | 14 | 12 | 12 | 12 |
| L75 | 25 | 21 | 21 | 20 |

Note: These statistics were calculated using the genome assemblies constructed using SPAdes and Satsuma2.

**Figure 2.**
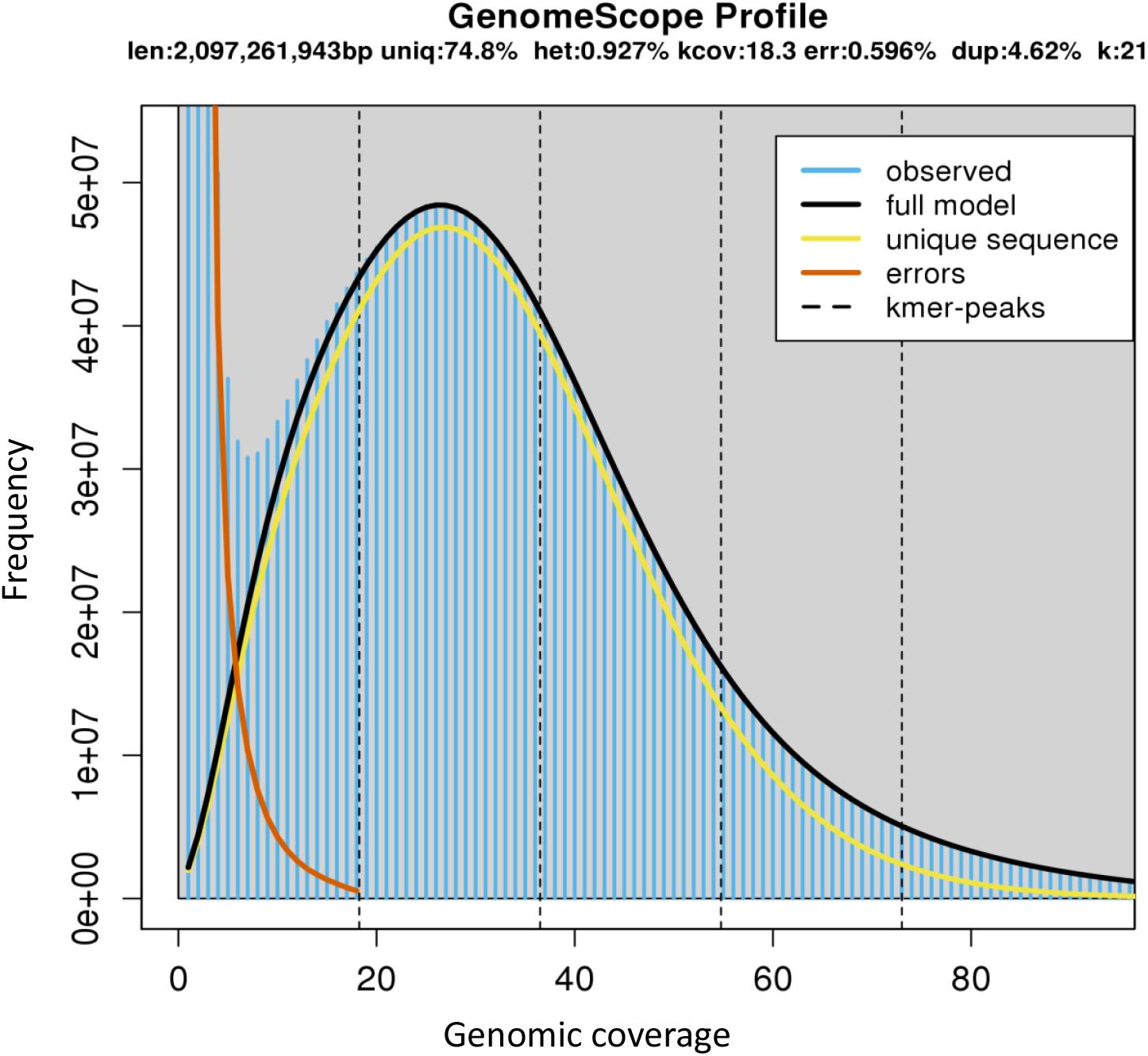
Genomic characteristics of Ongole breed of *B. indicus* species.

The forward and reverse amplicon sequences of COI gene were concatenated based on the aligned overlap to form a contiguous 655 bp sequence of the COI gene. Similarly, the forward and reverse sequences of 18S rRNA gene were concatenated to form complete 18S rRNA gene sequence of 1,756 bp. These resultant COI and 18S rRNA gene sequences were further aligned with the NCBI nr database using BLASTN. The COI gene showed 99.7% identity with *Bos indicus*, whereas the 18S rRNA gene showed 100% identity with the “Predicted: *Bos indicus* x *Bos taurus* 18S ribosomal RNA (XR_003508809.1)” sequence perhaps due to the absence of an experimentally verified 18S rRNA gene of *Bos indicus* in NCBI database. This assembled 18S rRNA gene was aligned to *Bos taurus* 18S rRNA gene (Accession: NR_036642.1) with 99.09% identity. Further, BLASTN alignment of each 18S rRNA partial sequence was performed against the contigs level assembly of each breed to obtain the 18S rRNA sequence containing contig. This helped to further extend the 18S rRNA sequence at both the ends using the reference contig sequence. This concatenated 18S rRNA sequence was aligned using BLASTN against the *Bos taurus* 18S rRNA sequence (NR_036642.1) to construct the full-length sequence. The obtained 18S rRNA gene sequence was found to be 1,869 bp long for Kasargod Dwarf, Kasargod Dwarf and Vechur, and 1,530 bp for Ongole.

### Exon-intron structure of genes responsible for adaptive traits of *B. indicus*

The exon-intron structures of 15 genes that are known to be associated with various characteristics such as milk quality, metabolism and immune response in *B. indicus* as reported in the previous studies were compared with the exon-intron structures of *B. taurus* genes. The analysis revealed eight genes that had variations in exon and intron numbers. Selected genes (*CSN1S1, CSN2*, and *PTGFR*) involved in milk quality and production showed a lesser numbers of intron-exons in Ongole compared to *B. taurus*. A similar observation was made for four out of five metabolism and immune response-related genes showing variations in intron-exon numbers, except for gene *AHCY*.

## DISCUSSION

In this study, we performed the whole genome sequencing, and *de novo* and reference-based assembly of *B. indicus* dwarf breeds (Kasargod Dwarf, Kasargod Kapila, Vechur) along with Ongole. Thus, this is also the first effort to sequence the smallest cattle breeds in the world. Reference-based genome assemblies of these cattle breeds helped in the identification of distinct *B. taurus* genes that were more diverged in *B. indicus* in terms of sequence identity. Genome sequencing data of dwarf breeds, and non-dwarf breed (Ongole) were used to construct *de novo* assembly of *B. indicus* cattle (Beja-Pereira et al., 2003). This cattle species showed moderate level of heterozygosity (0.927%), which is a limiting factor in generating a higher contiguous genome assembly (Asalone et al., 2020). However, despite this limitation, we were able to construct good quality *de novo* assemblies in terms of N50 values after scaffolding using Satsuma2, similar to other reported genomic studies (Dong et al., 2021). The gene (exon-intron) structures of 15 genes involved in the signature adaptive characteristics such as quality of milk, stature, stress tolerance, immune response, and metabolic pathways in *B. indicus* (Pryce et al., 2011; Dixit et al., 2020; Dixit et al., 2021; Elayadeth-Meethal et al., 2021), showed structural variation in exon-intron numbers between *B. taurus* and *B. indicus* Ongole breed. Seven of the genes in Ongole showed fewer exons than *B. taurus* among which *PFKP*, and *GPX4* (involved in glycolysis regulation and immune response, respectively (Dixit et al., 2020)) have a substantially higher difference (≥ 5 exons in *B. taurus*). However, *AHCY* involved in cellular processes such as epigenetic regulation (Vizán et al., 2021) showed a higher number of exons in Ongole compared to *B. taurus*. The resulting difference could be the result of inter-species variation in gene structure, which might influence the level of expression of these genes in the two *Bos* species (Seoighe et al., 2005; Yang, 2009). Such genes might also be responsible for the inter-species phenotypic adaptabilities of the cattle species, and provide valuable leads for further research. Thus, the availability of *de novo* genome assembly of *B. indicus* and 18S rRNA gene sequences of *B. indicus* breeds, and the knowledge of genes that showed structural variations between *B. taurus* and *B. indicus* will serve as a valuable resource for future studies on these indigenous cattle breeds.

## ACKNOWLEDGEMENTS

AC, MSB, and SM thank Council of Scientific and Industrial Research (CSIR), and RS thanks DST-INSPIRE for their research fellowships. We also thank NGS facility at IISER Bhopal, and the intramural research fund provided by IISER Bhopal. We thank Mr. P K Lal, Director of “Kasaragod Dwarf Conservation Society (KDCS)” for help in sample collection from the Kapila Gaushala, Ambalathara, Kanhangad, Kasaragod, Kerala, India.

## AUTHORS’ CONTRIBUTION

VKS conceived and coordinated the project. JP performed the morphological identification of *Bos indicus* breeds from their respective sample collection areas, and collected the samples. RS and SM performed DNA extraction, and Illumina sequencing. AC and VKS designed the computational framework of this study. MSB and AC performed the computational analyses, and constructed figures. SM and MSB constructed the 18S rRNA gene sequences. AC, MSB, and VKS interpreted the results. AC, MSB, SM, RS, and VKS wrote the manuscript. All authors have contributed and approved the final version of the manuscript.

## DISCLOSURES

The authors declare no conflict of interests.

## Notes

### Competing Interest Statement

The authors have declared no competing interest.

